# The learning curve of murine subretinal injection among clinically trained ophthalmic surgeons

**DOI:** 10.1101/2021.11.05.467533

**Authors:** Peirong Huang, Siddharth Narendran, Felipe Pereira, Shinichi Fukuda, Yosuke Nagasaka, Ivana Apicella, Praveen Yerramothu, Kenneth M. Marion, Xiaoyu Cai, Srinivas R. Sadda, Bradley D. Gelfand, Jayakrishna Ambati

**Author notes:** Correspondence: Jayakrishna Ambati, University of Virginia, Charlottesville, Virginia 22908. **Author contributions:** Conceptualization, P.H. and J.A.; Investigation, P.H., F.P., S.N., Y.N. and S.F.; Writing, P.H., F.P., S.N., K.M.M. with assistance from I.A., P.Y, X.C., S.R.S., B.D.G., and J.A. All authors had the opportunity to discuss the results and comment on the manuscript.

## Abstract

**Purpose:** Subretinal injection (SRI) in mice is widely used in retinal research, yet the learning curve (LC) of this surgically challenging technique is unknown.

**Methods:** To evaluate the LC for SRI in a murine model, we analyzed training data from 3 clinically trained ophthalmic surgeons from 2018 to 2020. Successful SRI was defined as either the absence of retinal pigment epithelium (RPE) degeneration after phosphate buffer saline injection and the presence of RPE degeneration after *Alu* RNA injection. Multivariable survival-time regression models were used to evaluate the association between surgeon experience and success rate, with adjustment for injection agents, and to calculate an approximate case number to achieve a 95% success rate. A Cumulative Sum (CUSUM) analysis was performed and plotted individually to monitor each surgeon’s simultaneous performance.

**Results:** Despite prior microsurgery experience, the combined average success rate of the first 50 cases in mice was only 27%. The predicted SRI success rate did not reach a plateau above 95% until approximately 364 prior cases. Using the 364-training case as a “cutoff” point, the predicted probability of success before and after the 364^th^ case was 65.38% and 99.32%, respectively (*P* < 0.0001). CUSUM analysis showed an initial upward slope and then remained within the decision intervals with an acceptable success rate set at 95% in the late stage.

**Conclusions:** This study demonstrates the complexity and substantial LC for successful SRI in mice with high confidence. A systematic training system could improve the reliability and reproducibility of SRI-related experiments and improve the interpretation of experimental results using this technique.

**Translational Relevance:** Our prediction model and monitor system allow objective quantification of technical proficiency in the field of subretinal drug delivery and gene therapy for the first time.

## INTRODUCTION

Subretinal injection (SRI) is a fundamental technique for accessing and delivering agents into the subretinal space. Clinically, SRI is used in vitrectomy surgeries to treat submacular hemorrhages^1^ and retinal folds,^2^ as well as to deliver gene therapy to treat retinal dystrophies.^3^ In retinal research, SRI plays a key role in elucidating the pathogenesis of disease through its use in rodent models for gene and cell therapy.^4-6^

Three techniques are commonly employed to access the subretinal space during SRI. The transcorneal route, which involves the needle bypassing the iris and lens through the pupil, is often utilized in studies where induction of total retinal detachment is desirable and maintenance of clear media post-operatively is not required.^7^ The transscleral posterior route, which enables access to the subretinal space without entering the vitreous cavity or penetrating the retina, is the mainstay procedure in newborn mice.^8, 9^ Finally, the pars plana route, in which the needle is inserted via a sclerotomy, is utilized when precise, localized areas of subretinal injection is desired as it permits direct visualization of the trajectory of the needle within the vitreous as well as of the exact site of injection.^10^

A paramount concern in animal models employing SRI is the accurate interpretation of the retinal and RPE phenotypes during the follow-up period. Minimizing the manipulation of the eye and controlling the position of the needle at all times under direct visualization are two critical advantages of the pars plana route. However, the pars plana route can be challenging to master. The mouse eye has a vitreous volume of less than 10 µL and its lens occupies three-quarters of the posterior segment,^11^ imposing technical challenges even for experienced surgeons. There are no reported data for the training time required to master SRI in mice. This is a crucial gap in knowledge, particularly because of increasingly widespread studies employing SRI in mice.

Therefore, we sought to establish the temporal kinetics of attaining proficiency in SRI in mice to assess the outcomes of experiments relying on this technique with confidence. We employed the learning curve (LC) methodology, which has been widely adopted to describe the rate of progress in learning a new technique over time. In clinical surgery, LC analysis is used to determine the number of procedures a surgeon needs to perform before achieving consistently good outcomes.^12^ The use of the LC to improve health care outcomes has a parallel in experimental research, wherein proper training before conducting formal experiments with unknown outcomes can minimize misinterpretation of results and misleading conclusions. This study aimed to define and interpret the success or failure of SRI in mice, evaluate the learning curve of trained ophthalmologists to master the SRI technique using the pars plana route, and implement an audit tool called Cumulative Sum (CUSUM) to monitor the learning process.

## Methods

### Animals and Materials

Wild-type C57BL/6J mice between 6-8 weeks of age (The Jackson Laboratory, Bar Harbor, ME, USA) were used in this study. The mice were anesthetized by 100 mg/kg ketamine hydrochloride (Fort Dodge Animal Health, Overland Park, KS, USA) and 10 mg/kg xylazine (Akorn, Lake Forrest, IL, USA). Animals were placed on a heating pad at 37 °C and pupils were dilated using topical 1% tropicamide and 2.5% phenylephrine hydrochloride (Alcon, Fort Worth, TX, USA). All animal experiments were approved by the University of Virginia Institutional Animal Care and Use Committee and performed according to the ARVO Statement for the Use of Animals in Ophthalmic and Visual Research.

For this study, the surgeons performed SRI of 1x phosphate buffer saline (PBS), an inert substance, or *Alu* RNA, which induces RPE degeneration.^13-21^ Under a biosafety cabinet, we diluted 10x PBS ultrapure-grade (VWR, Radnor, PA, USA) to 1x PBS. 300 ng *Alu* RNA was prepared as previously described and injected.^13, 19^ AAV2-hRPE (0.8)-iCre-WPRE and AAV2-CMV-null were obtained from Vector Biolabs.

### Subretinal injection technique

Eyes were anesthetized using topical 0.5% proparacaine (Bausch & Lomb, Rochester, NY, USA) and topical methylcellulose 2% (Akorn, Lake Forrest, IL, USA) was quickly applied to avoid dry eye-related media opacities.^22^ A sclerotomy was placed 0.5-1.0 mm from the limbus using a 30-gauge needle. A custom-made optical lens was placed over the cornea to permit a clear view of the posterior retina under a surgical microscope (Leica Microsystems, Wetzlar, Germany). A 37-gauge needle attached to a 5 µL syringe (Ito Corporation, Tokyo, Japan) was introduced through the sclerotomy. Under direct visualization, 0.5 µL of 1×PBS or *Alu* RNA was injected through a posterior retinectomy until a small restricted bubble was visualized, ensuring proper injection. At the end of the procedure, eyes were covered with antibiotic ointment (Bausch & Lomb, Rochester, NY, USA).

### Outcomes and retinal imaging

Seven days after the injection, pupils were dilated and fundus imaging was performed with a TRC-50 IX camera (Topcon, Tokyo, Japan) linked to a digital imaging system (Sony). A successful injection was defined by the absence of intraoperative complications (lens touch, diffuse retinal detachment, or retinal bleeding) combined with absence of degenerative changes after PBS injection or presence of degenerative changes after *Alu* RNA injection. Immunofluorescence staining of zonula occludens-1 (ZO-1) on retinal pigment epithelium (RPE) flat mounts was used to ascertain the presence or absence of degeneration.

### Immunofluorescence

After enucleation, eyes cups and retina were carefully removed. RPE flat mounts were fixed in 2% paraformaldehyde, stained with rabbit polyclonal antibodies against mouse ZO-1 (1:100, Invitrogen, Carlsbad, CA, USA) and visualized with Alexa Fluor 594 (Invitrogen). For immunofluorescence staining for Cre expression, fresh, unfixed mouse eyes were embedded in Optimal Cutting Temperature Compound (Fisher), frozen in isopentane precooled by liquid nitrogen and then cryosectioned. A rabbit polyclonal to Cre recombinase was used (1:50; Abcam). Using a confocal microscope (Nikon A1R confocal microscope system), flat mounts were imaged and assessed by a blinded operator. Degeneration in RPE flat mounts was defined using an established protocol.^19^

### Study cohort and data source

The experience of three trained ophthalmologists with SRI in mice using the pars plana route was recorded between 2018 and 2020. We computed the number of surgeries, success rate and complication rate for all surgeons.

### Multivariable models

Surgeon’s experience was coded as the number of subretinal injection eyes performed by the surgeon prior to the index case. The association between injection outcomes and surgeon’s experience was tested, correcting for the type of injection (PBS or *Alu* RNA), using a survival regression model clustered by the surgeon. To evaluate the association between a surgeon’s experience and the results of the injection observed on day 7, we employed a multivariable, parametric survival time regression model with a log-logistic distribution to model the probability of freedom from failure against surgeon experience, as reported by Vickers et al.^23, 24^ The number of cases needed to reach a 95% success rate was defined as the “cutoff”. Proportion comparisons between the groups before the “cutoff” point and afterward were performed with a Fisher exact test, and numerical comparisons were conducted with a Mann-Whitney U test. The level of statistical significance was defined as *P* < 0.05.

### Cumulative Sum (CUSUM) analysis

The CUSUM score was determined by the formula: CUSUM C_n_ =max (0, C_*n*-1_ + X_*n*_ -k), where C = case, n = number of injections since the start of training, X_n_=0 (success), X_n_ =1 (failure), and k = reference value that is determined by the acceptable and unacceptable failure rate.^25^ A narrower decision interval is easier to cross, which allows less chance of consecutive errors. The tradeoff for a narrow decision interval is the excess of false alarms in loss of performance. For visualization, the number of cases was plotted on the x-axis, and the CUSUM score on the y-axis.

### The learning curve of complication rates

To solve the learning curve for a dichotomous outcome such as complication rates, cumulative complication numbers were plotted against case numbers. Cataract, bleeding and other complications were included in the cumulative complication number in keeping with prior reports of learning curves^26^. The average complication rate was the total number of relevant complications over the entire case series. The derivatives of the association curves fitted to the overall data sets were calculated, and the corresponding case numbers were regarded as “cutoff” points. Proportional comparisons between the groups before the “cutoff” points and afterward were performed with Fisher’s exact tests, and numerical comparisons were conducted with a Mann-Whitney U tests. Statistical significance was defined as *P* < 0.05.

## RESULTS

### The learning curve for SRI in mice

We used PBS as the negative control and *Alu* RNA as the positive control for RPE degeneration to assess the technical proficiency in the training period. Based on our prior studies,^13-21^ successful SRI of PBS does not result in RPE degeneration after 7 days. Successful subretinal delivery of *Alu* RNA results in RPE degeneration evident after 7 days.^13, 14, 16-19, 27^

Prior to commencing subretinal injection training, all three trainees had completed three-year surgical residency programs in ophthalmology with experience in microsurgeries in people. During the training period, surgeons 1, 2 and 3 injected 455, 365 and 415 eyes, respectively, using the pars plana technique. Despite extensive prior experience in clinical ophthalmology, the success rate for the first 50 cases for each surgeon was under 40%, with a combined average success rate of 27%. From cases 50 to 350, each surgeon consistently improved over time, until attaining an average 90% success rate between cases 301 - 350 (Figure 1).

**Figure 1.**
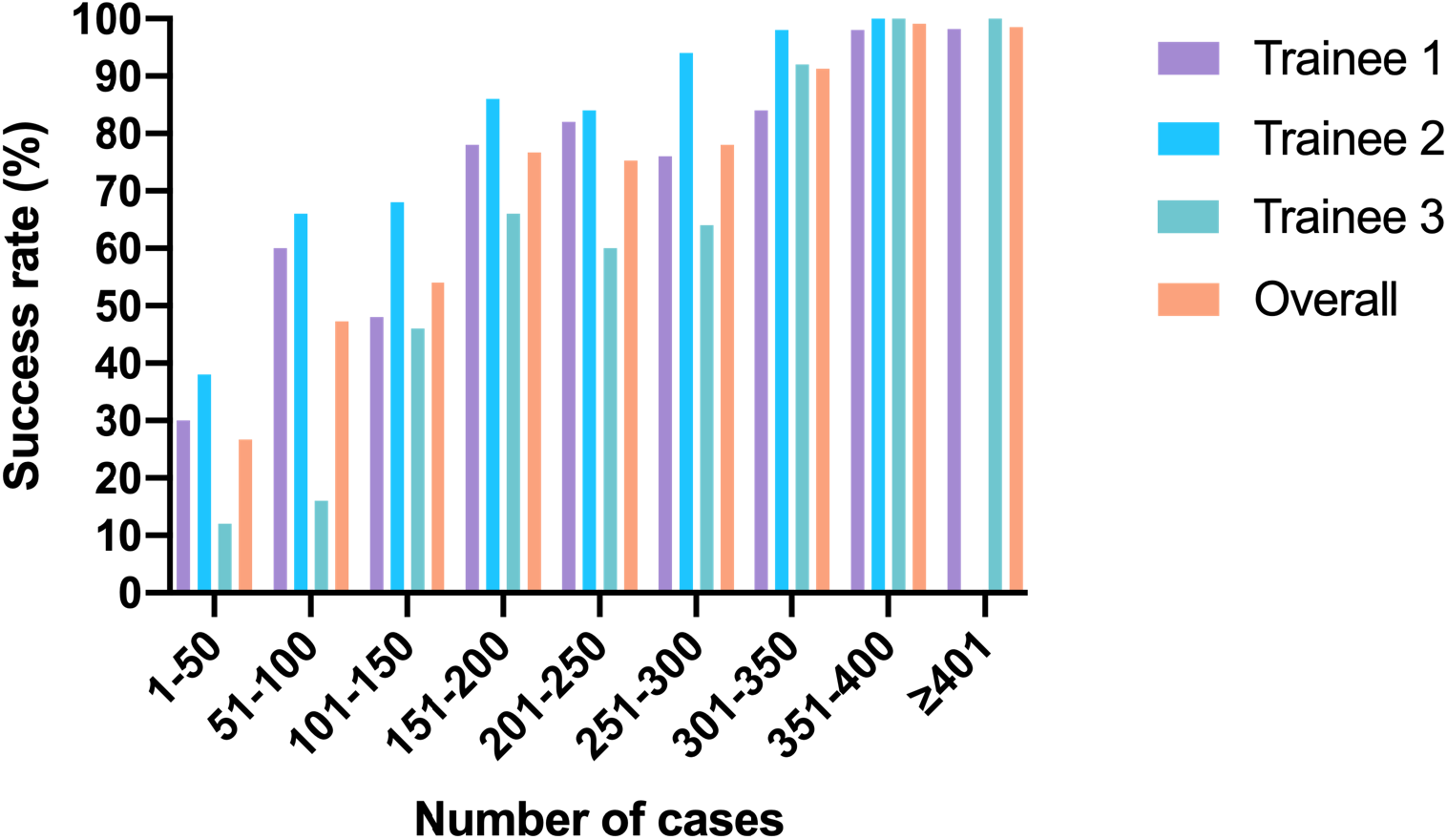
Descriptive analysis of each surgeon’s experience and success rate.

Using data from all three surgeons, we built a predictive model to estimate the number of cases necessary to achieve a 95% success rate (Figure 2). After correcting for the type of SRI (PBS or *Alu* RNA), the model indicated an accelerated improvement rate up until 150 cases, followed by a phase of slower improvement with higher variability from 150 to 250 cases. This slower improvement phase coincided with the introduction of SRI of *Alu* RNA into the training. After 250 cases, the curve ascended with at a steeper slope, reaching a 95% success rate at 364 cases. Using the 364^th^ case as the “cutoff” point, the predicted probability of success before and after the 364^th^ case was 65.38% and 99.32%, respectively (*P* < 0.001).

**Figure 2.**
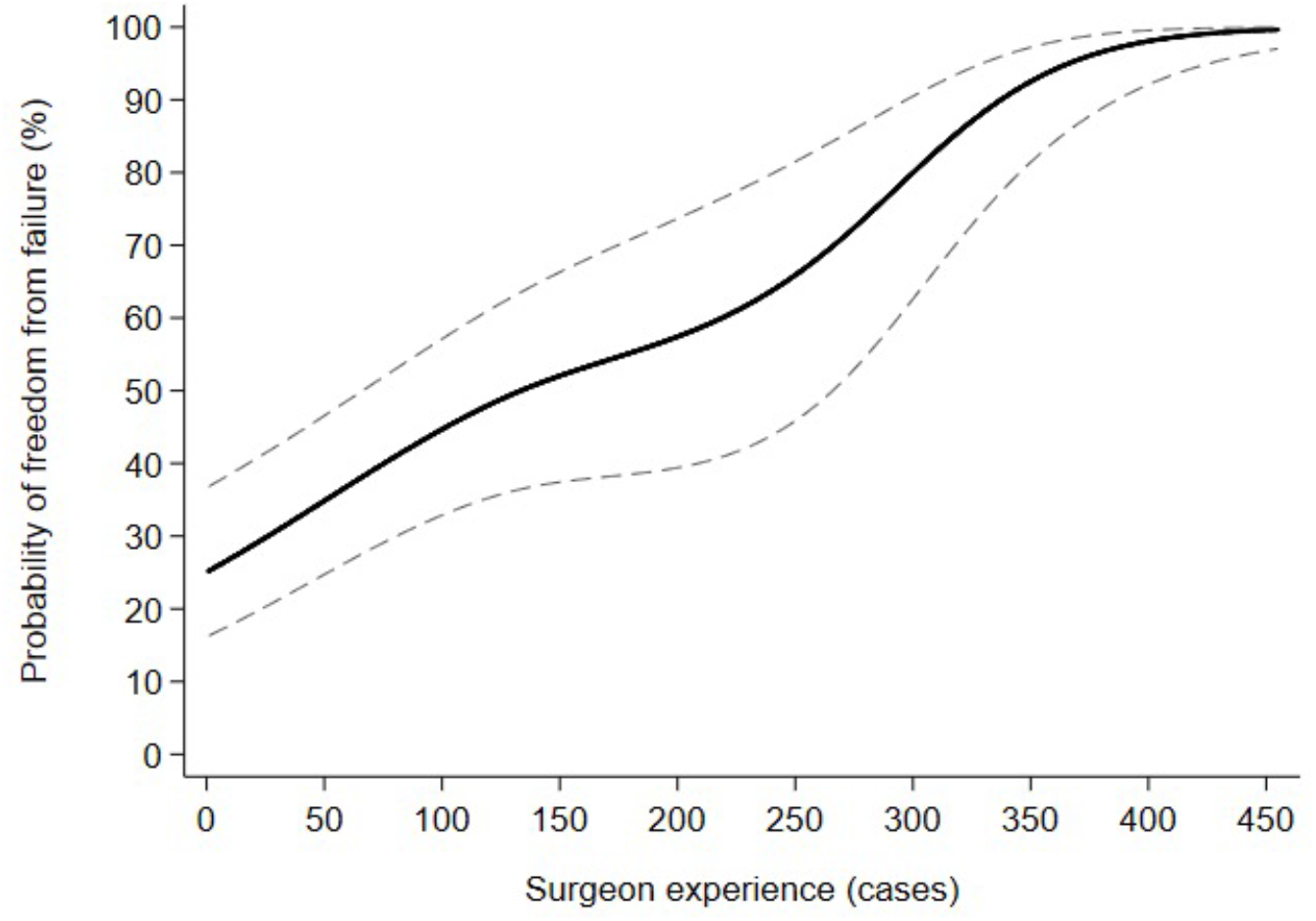
Predictive surgical learning curve for successful subretinal injection in mice. Predicted probability (black central curve) and 95% confidence intervals (dashed lines) were plotted against increasing surgeon experience. The injection agents (PBS or *Alu* RNA) were interpreted as covariates to adjust the regression model.

### CUSUM analysis of SRI performance

CUSUM analysis enables early quantitative detection of changes in task performance.^25, 28^ In the clinical application of CUSUM, when a surgeon’s performance is inside the range of a prespecified acceptable success rate, the CUSUM score decreases, and the CUSUM curve becomes flat or turns downwards. Conversely, the graph turns upwards when a significant reduction in performance occurs. If the curve continues rising and crosses a decision interval (h), this indicates that the performance is not acceptable at that set level. We applied these functions of CUSUM to monitor the performance of our trainees. In our analysis, the decision interval (h) was calculated with the unacceptable rate of failure set at higher than 5%. Other parameters used to calculate the CUSUM are displayed in Table 1.

**Table 1.** CUSUM charting design for monitor training.

| Specifications | Parameters |
| --- | --- |
| Acceptable rate of failure for subretinal injection, $\pi_1$ | 0.001% |
| Unacceptable rate of failure for subretinal injection, $\pi_2$ | 5% |
| Reference value k, calculated based on $\pi_1$ and $\pi_2$ using methods described here<sup>46</sup> | 0.0082 |
| *h, †IC-ARL and ‡OC-ARL, calculated based on k | h=1.00<br>IC-ARL=10000<br>OC-ARL=20 |

\*h: decision interval
†IC-ARL: in-control average run length
‡OC-ARL: out-of control average run length

The CUSUM curves for a successful outcome after SRI of all three surgeons during the entire training period and a detailed chart of the last 150 cases are displayed in Figure 3. Due to the very low threshold for the unacceptable rate of failure (5%) and the resulting small decision interval, all surgeons experienced an increase in the CUSUM score, with the curve sloping upwards during the beginning of the training. Surgeon 1 was able to maintain the same decision interval after 355 cases. Surgeon 2 reached the desirable success rate after 265 cases and surgeon 3, after 315 cases. Surgeons 1 and 2 had ascending points at cases 401 and 304, respectively. However, as the failures were far apart, they did not result in the graph crossing the decision interval.

**Figure 3.**
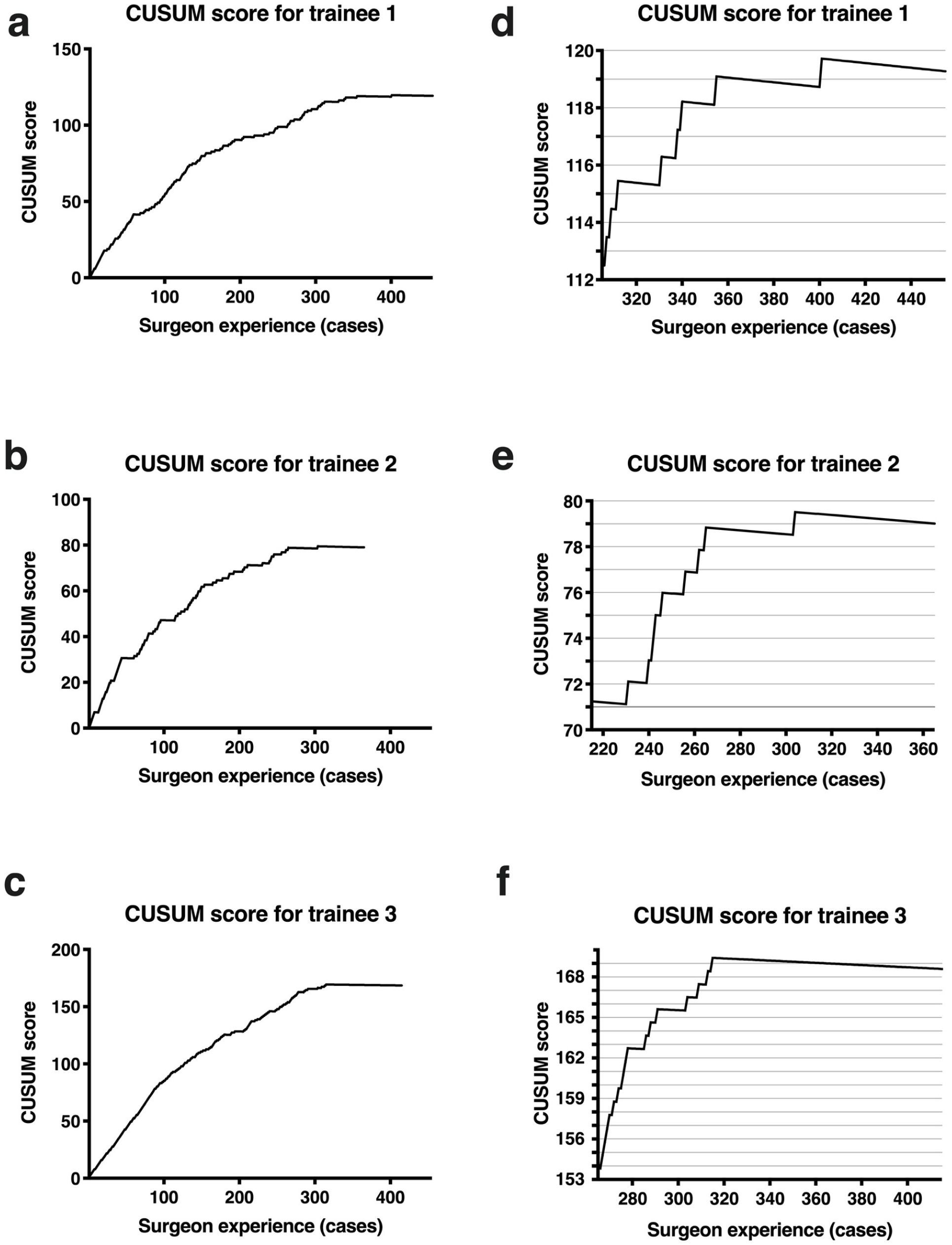
CUSUM chart as an auditing simultaneous monitoring tool. (a - c) CUSUM chart of each surgeon for the entire training period based on an acceptable success rate set at 95%. (d - f) Magnified CUSUM chart of the last 150 cases of each surgeon. The gray horizontal grids in (d - f) indicate the decision interval (h) that is equal to 1.

### Complication rates

Cataract formation and bleeding were the most common complications in our SRI training records. When these complications occur, they can seriously interfere with outcome assessment. Using these metrics, we analyzed two surgeons’ complication rates classified by cataract, bleeding, and others. The rates of complications decreased with increasing surgeon experience. For surgeon 1, a total of 75 (16.48%) complications (16 cataract cases, 24 bleeding cases and 35 other complications) occurred over 455 cases. Graphs of the cumulative number of complications over time display a sharp plateau starting at case 136, 141, and 155 for cataract, bleeding and other complications, respectively. At these points, the derivatives of the association curves equations equal the overall complication rates of 3.51% (cataract cases), 5.27% (bleeding cases), and 7.69% (other complications), respectively. Fourteen cataract cases, 20 bleeding cases, and 28 cases with other complications occurred before the plateau cases (10.29%, 14.18%, 18.06%) compared to only 2, 4, and 7, respectively, after the plateaus (0.63%, 1.27%, 2.33%). Therefore, 87.5% of cataract, 83.33% of bleeding, and 80% of other complications occurred before the plateau cases and these proportions were statistically significant, where Fisher’s Exact Test p-values for cataract cases, bleeding cases and other complications were 1.83 × 10^−6^, 7.67 × 10^−8^, and 8.66 × 10^−9^, respectively. For the second trainee, a total of 55 (15.07%) complications (17 cataract cases, 8 bleeding cases and 30 other complications) occurred within 365 cases. Graphs of the cumulative number of complications over time display a sharp plateau starting at case 107, 111 and 130, which are where the derivative of the association curve equations equal the overall complication rates of 4.66% (cataract cases), 2.19% (bleeding cases), and 8.22% (other complications), respectively. Fourteen cataract cases, 7 bleeding cases, and 21 cases with other complications occurred before plateau cases (13.08%, 6.31%, 16.15%) compared to only 3, 1, 9 after the plateau (1.16%, 0.39%, 3.83%). Therefore, 82.35% of cataract, 87.5% of bleeding, and 70% of other complications occurred before plateau cases and these proportions were statistically significant, where Fisher’s Exact Test p-values for cataract cases, bleeding cases and other complications were 5.31× 10^−6^, 1.25× 10^−3^, and 8.62 × 10^−5^, respectively. Both of the surgeons’ instantaneous complication rates decreased, plotted in the lower panel of Figure 4.

**Figure 4.**
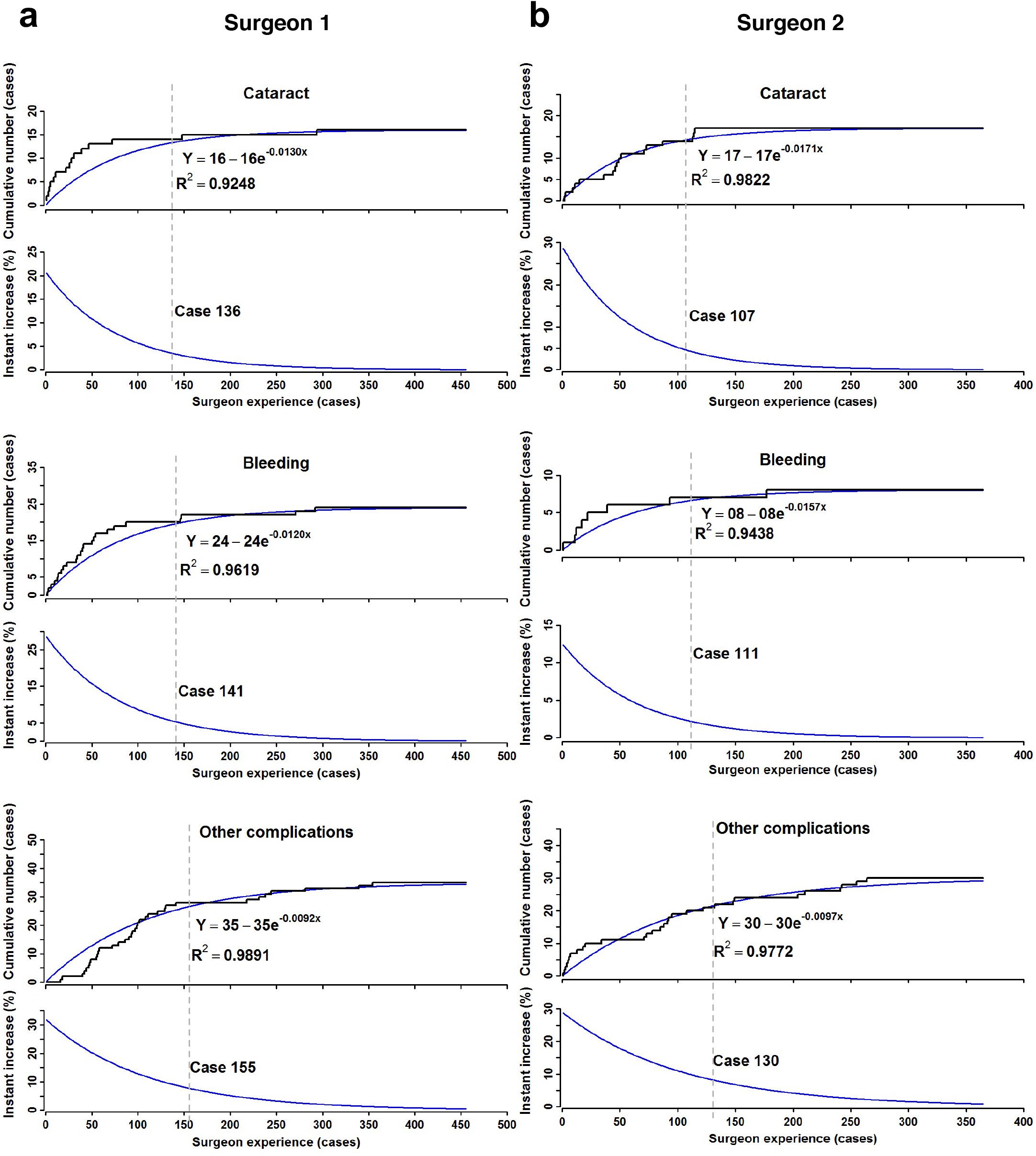
Cumulative number of complications. Cumulative number of complications (cataract, bleeding and other complications) were plotted against surgeon 1 (a) and 2’s (b) training case numbers. Association curves were generated to fit the plots. Derivatives of the equations solved for complication rates identify the learning curves at case 136, 141, 155 (surgeon 1) and 107, 111, 130 (surgeon 2) for cataract, bleeding and others, respectively. Both of the trainees’ instant complication rate decreased with increasing experience.

## DISCUSSION

In this study, we quantified the success or failure of three trained ophthalmic surgeons learning SRI in mice. Based on our findings, we predict that a trained ophthalmologist needs to perform an average of 364 SRI procedures in mice to achieve a 95% success rate. Observationally, these findings are in line with our roughly 20-year history of supervising clinically-trained surgeons learning this technique. Similarly, we estimate that trainees lacking similar extensive prior ophthalmic surgery experience would require a significantly higher number of cases to achieve proficiency. We recommend using a quantitative tool such as CUSUM to assess the surgeon’s competency in real time.

An important aspect to consider when assessing the LC of a surgical procedure is the variability measured that defines success. When defining “success” for an LC analysis, there are typically two options: (1) intraoperative success or (2) positive outcome of the procedure.^12^ The optimal choice of a variable will depend on the type of procedure. For instance, a bullous retinal detachment without intraoperative complications may constitute evidence that the technique was successfully employed to deliver the agent to the subretinal space. However, this does not imply that the procedure was not harmful to the eye. Our experience shows that an extensive retinal detachment following SRI, comparable to those described in the former studies ^29, 30^, invariably leads to RPE and retinal degeneration. Conversely, proper delivery of PBS should not induce RPE degeneration after successful training. Hence, setting positive and negative outcomes for SRI in the training period is important to identify and rectify potential errors in this multi-step complex operation.

Low reproducibility rates can plague preclinical studies, with reports that cumulative (total) prevalence of irreproducible preclinical research exceeds 50% (ranging from 51 - 89%),^31-35^ resulting in around $28 billion wasted annually in the United States.^36^ Lack of training, along with scientific culture and incentives, have all been blamed for irreproducibility.^37^ Our study reinforces the importance of an extensive training program in SRI in mice to increase confidence in results using from this technique.

Irrespective of the SRI technique used, some studies have reported morphological and functional alterations after SRI of PBS in mice.^29, 30^ However, in those studies, the authors clearly describe or present images of extensive retinal detachment that induce iatrogenic damage, as a successful procedure. In contrast, after our training periods, the rate at which SRI of PBS does not induce RPE degeneration at day 7 confirmed by fundus photo and ZO-1 staining, exceeds 95%. Indeed, other studies have also demonstrated that SRI of PBS in mice is safe.^38, 39^

In our experience, even if the total injected volume of PBS is the same, the extent of retinal detachment (shape, height and range) can vary significantly when visualized by microscopy, due to technical parameters. We have found that to perform a successful SRI, the injection speed should be slow, allowing the liquid to gradually open the subretinal space, and the force applied on the syringe should be gentle, avoiding a sudden fluctuation of the intraocular pressure. Improper preparation of PBS (level of purity, sterilization, osmotic pressure, etc.) also could also lead to an artifactual degeneration of RPE cells. Our group has successfully used the technique described here to identify new molecular mechanisms driving the pathogenesis of age-related macular degeneration and propose new therapeutic targets.^13-21^ This highlights the importance of the training program for SRI in mice and the importance of identifying an appropriate outcome to be evaluated during the follow-up period.

The regression model reported in this study shows that experienced surgeons need to perform approximately 364 cases of SRI in mice to reach the desirable success rate of 95%. Such a high bar for success is essential to ensure reliable and consistent results in this procedure. In comparison, three second-year ophthalmology residents took an average of 38 cases to reach an acceptable rate of success in phacoemulsification in people,^40^ highlighting the relative complexity of the SRI procedure in mice.

In rats, SRI also demands fewer cases to achieve proficiency.^41^ Several factors can contribute to this significant difference in the LC. First, the mouse eye, with an approximate axial length of only 3 mm^42, 43^, is significantly smaller than that of rat. Simultaneously, the lens diameter in mice is proportionally larger (2 mm)^44^. The limited residual space for manipulation in mice increases the complexity of the intraocular operation. Second, the definition of success described in the rat study^41^ took into account only intraoperative aspects and its related complications, without assessing the anatomical or functional consequences of the procedure.

CUSUM charts have been widely used as a method for real-time performance monitoring in surgical procedures.^25, 45^ We introduced CUSUM charts into the SRI training program to track the trainee’s performance and provide objective, quantitative feedback based on desired standards. Its graphic display is advantageous in its ability to be easily understood and its reflection of performance quality. If the CUSUM curve shows a continued upward trend, more guidance, feedback, and attention should be given to the trainee by senior researchers to help improve the technique. Conversely, when the CUSUM curve remains steady within the decision intervals, it indicates the trainee has achieved proficiency. In summary, CUSUM can be applied both during and after training to ensure reproducibility and validity of the experimental results.

Our study has several strengths. First, we generated a probability model addressing the relationship between surgeon experience and experimental competency beyond merely comparing the first and the last dozen cases as a readout in the training periods. Second, we used data from three different surgeons with similar prior experience in microsurgery. Third, we defined success by the presence or absence of RPE degeneration after 7 days from the injection by fundus image and immunofluorescence staining of RPE flat mounts, which, compared to restricting the definition of success to intraoperative readouts, better reflects the actual effect of this procedure on the RPE. Fourth, using SRI of both positive (*Alu* RNA) and negative (PBS) controls increases confidence that successful training reflects true competency in this technique. Fifth, this study is the first application of the CUSUM score to quantify skill training in the basic sciences for improving reproducibility. Limitations of this study include the relatively small number of trainees assessed. Since all trainees had previous experience in microsurgery in the clinic before the beginning of the training period, the number of cases to master this technique may differ significantly for a person without such clinical surgical experience.

## Acknowledgments

J.A. has received support from NIH grants (R01EY028027, R01EY29799, R01EY031039), the DuPont Guerry, III, Professorship, and a gift from Mr. and Mrs. Eli W. Tullis; B.D.G. has received support from NIH grants (R01EY028027, R01EY031039, R01EY032512), BrightFocus Foundation, and the Owens Family Foundation.

J.A. has received support from the UVA Strategic Investment Fund, NIH grants (R01EY028027, R01EY29799, R01EY31039), DuPont Guerry, III, Professorship, and a gift from Mr. and Mrs. Eli W. Tullis; B.D.G. has received support from NIH grants (R01EY028027, R01EY031039, R01EY032512), BrightFocus Foundation, and the Owens Family Foundation.

## Competing interests

J.A. is a co-founder of iVeena Holdings, iVeena Delivery Systems, and Inflammasome Therapeutics, and has been a consultant for Allergan, Boehringer-Ingelheim, Immunovant, Olix Pharmaceuticals, Retinal Solutions, and Saksin LifeSciences unrelated to this work. S.R.S. has been a consultant for 4DMT, Abbvie/Allergan, Apellis, Amgen, Centervue, Heidelberg, Iveric, Novartis, Optos, Oxurion, Regeneron, and Roche/Genentech, received speaker fees from Novartis, Nidek, Carl Zeiss Meditec, and Optos, and received research instruments from Carl Zeiss Meditec, Nidek, and Topcon, Centervue, Optos, Heidelberg unrelated to this work; J.A. and B.D.G. are co-founders of DiceRx. J.A., S.N., I.A., F.P., and B.D.G. are named as inventors on patent applications on macular degeneration filed by their university.

